# Microbiome Dynamics in Tank- and Pond-Reared GIFT Tilapia

**DOI:** 10.1101/2024.11.16.622739

**Authors:** Jérôme Delamare-Deboutteville, Mahirah Mahmuddin, Han Ming Gan, Charles Rodde, Laura Khor, David Verner-Jeffreys, Vishnumurthy Mohan Chadag, John A.H. Benzie

**Affiliations:** WorldFish, Aquatic Food Biosciences, Jalan Batu Maung, Batu Maung, Penang 11960, Malaysia; Patriot Biotech Sdn Bhd, Bandar Sunway, 475000 Subang Jaya, Selangor, Malaysia

**Author notes:** **Correspondance:** Jérôme Delamare-Deboutteville.

## Abstract

Tilapia (*Oreochromis* spp.) are among the most widely cultivated freshwater finfish species worldwide. The industry increasingly relies on tilapia strains selected for improved growth and other traits, particularly the Genetically Improved Farmed Tilapia (GIFT) strain. Despite the industry’s reliance on tilapia, knowledge of microbiome dynamics in reared tilapia remains limited. Understanding normal successional patterns in the microbiome of farmed tilapia is essential to identify characteristics of what constitutes a healthy microbial community. In this study, we assessed the microbiomes of tank and pond-reared GIFT tilapia by analyzing 568 samples, including water, gut, skin, and gill microbiomes of tilapia, from tank systems housing the source GIFT populations in Malaysia, and compared them to those reared in earthen ponds on another farm in Malaysia. A total of 2,307 amplicon sequence variants (ASVs) were identified, encompassing a broad taxonomic diversity of 39 phyla, 86 classes, 180 orders, 299 families, 501 genera, and 399 species.

Our findings elucidated distinct microbial community structures between rearing environments and across fish tissues, shedding light on intricate host-microbe interactions shaped by environmental conditions and management practices. The gut microbiome of tank-reared tilapia was dominated by Fusobacteriota (71.14%), in contrast to pond-reared fish (22%), while other taxa, such as Bacteroidota, Firmicutes_A, and Cyanobacteria, also varied markedly between environments and sampling periods. Skin and gill samples showed notable variability in the relative abundances of Fusobacteriota and Deinococcota between the two rearing sites. Principal Coordinates Analysis (PCoA) highlighted the distinct clustering of samples by rearing environment, particularly within gut microbiomes. Biomarkers such as Cyanobiaceae (pond water) and Sphingomonadaceae (tank water) underscored the impact of rearing conditions on microbial composition.

These results establish valuable baseline information on the types of bacteria associated with healthy, genetically defined (GIFT) tilapia strains. This foundational information will help identify specific microbial taxa associated with beneficial or detrimental effects on tilapia health and productivity across varying rearing conditions. Such insights can guide the development of targeted microbiome management strategies to enhance tilapia health and optimize performance.

## Introduction

Aquaculture is a vital solution to meet the escalating global demand for seafood commodities amidst declining wild fish stock populations and mounting food security challenges of an ever-expanding global population (FAO, 2024; Tacon and Shumway, 2024). Among the diverse array of species cultivated in aquaculture systems, tilapia (*Oreochromis* spp.) is the most widely farmed freshwater finfish, being produced in over 140 countries (Dong et al., 2023). Many species of tilapia are produced globally, dominated by Nile tilapia (*Oreochromis niloticus* L.), cultured mainly in lower-middle-income countries (LMICs) across the Southeast Asian, African, and South American continents (Debnath et al., 2023). Tilapia stands out for its adaptability to diverse farming conditions, rapid growth, resistance against disease, and widespread consumer acceptance, making it a cornerstone of worldwide aquaculture production.

The Genetically Improved Farmed Tilapia (GIFT) strain represents a significant advancement in tilapia aquaculture, characterized by its rapid growth, high productivity, and adaptability to various geographies and farming environments (Hamzah et al., 2014; Ponzoni et al., 2011). Developed through selective breeding programs, the GIFT strain is renowned for its superior performance traits, making it a preferred choice among aquaculturists worldwide. Originating from internationally collaborative efforts coordinated by the International Center for Living Aquatic Resources Management (ICLARM, now the WorldFish center) in cooperation with Norwegian and Philippine partners, the GIFT strain was first established in the 1980s through systematic selection for growth rate, survival, and other economically important traits (Ponzoni et al., 2011). Over the years, continuous breeding and selection have refined the GIFT strain, resulting in improved genetic lines capable of outperforming conventional tilapia strains in terms of growth efficiency and disease resistance (Barría et al., 2020; Hamzah et al., 2014). With the recent publication of the GIFT tilapia genome (Etherington et al., 2022), the characterization of the microbial systems (microbiomes) of both the host-farmed species and their aquatic environments becomes even more imperative. Characterizing the gills, skin outer mucosal surface, and gut microbiomes and their relationship to the host’s genetics can provide insights into the mechanisms underlying host-microbe interactions and their contributions to key performance traits such as growth rate, fish health, disease resistance, nutrient metabolism, and overall host physiology. Furthermore, by identifying specific microbial taxa associated with beneficial or detrimental effects on GIFT health and productivity, targeted interventions can be developed to modulate the microbiome and enhance aquaculture performance.

A number of investigations have been undertaken to elucidate the microbiome dynamics of Nile tilapia. (Wu et al., 2021) conducted a study examining alterations in the gut microbiota of GIFT strains cultivated in a commercial fish farm (cement ponds) located in Hubei Province, China, following inoculation with an inactivated bivalent *Aeromonas hydrophila/Aeromonas veronii* vaccine. Their findings revealed a significant reduction in the relative abundance of fish pathogens within the gut post-vaccination. Meanwhile, (Zhu et al., 2022) explored disparities in the gut microbiota in tilapia reared in both pond culture systems and in-pond raceway systems, highlighting a progressive divergence in gut microbiota as the specimens mature. Additionally, (Parata et al., 2020) investigated the influence of diet on the gut microbiome of GIFT strains reared in earthen ponds in Papua New Guinea. Their study demonstrated that dietary composition influences the proliferation of specific bacteria, particularly those associated with the breakdown and digestion of consumed carbohydrates. Moreover, fish fed a consistent diet tended to exhibit a more stable microbiome. In addition to investigations into the gut microbiome, recent studies have ventured into the skin microbiome of Nile tilapia cultivated in Malawi, aiming to establish connections between the skin microbiome and the microbiome of the rearing water (McMurtrie et al., 2022). These investigations underscored compositional distinctions between fish skin and water microbial communities yet revealed shared taxa. Notably, pond locations strongly influenced the water microbiome compared to the fish skin microbiome. Furthermore, the skin microbiome holds promise as a potential indicator for monitoring disease in fish populations. Further expanding on microbiome research, (Elsheshtawy et al., 2021) investigated the microbiomes of the gills, skin, and pond water of Nile tilapia co-cultured with grey mullet (*Mugil capito*) in semi-intensive polyculture fish farms in Egypt. This study showed distinct gills bacterial communities from those of the skin in both species. Gill microbial communities being clearly species specific, while skin communities showed some overlap between both species, with the rearing water exhibiting highest abundance and richness.

Despite the growing interest in microbiome research worldwide, studies specifically targeting the microbiome of Malaysia-derived GIFT tilapia strains that have been distributed worldwide are lacking. This knowledge gap is particularly noteworthy considering the unique environmental conditions and microbial diversity inherent to tropical ecosystems, which can significantly impact aquaculture dynamics and fish health. Consequently, this gap poses a significant obstacle to the development of tailored aquaculture management strategies adapted to the region’s distinctive environmental and microbial landscapes. The composition and dynamics of the tilapia microbiomes can vary significantly due to differences in farming and breeding practices and environmental factors such as water quality, nutrient availability, and sunlight exposure. Notably, in earthen ponds, which are expansive bodies of water, these ecosystems allow for the proliferation of microbial and algae communities with minimal water exchange. In addition, the abundance of sunlight throughout the year in tropical countries will foster the growth of algae and other phototrophic and eukaryotic organisms in these ponds, serving as an alternative nutrient source for tilapia. In contrast, tank water cultivation systems, characterized by their smaller scale and controlled environment, necessitate frequent water changes and rigorous management practices. These systems afford greater control over water quality parameters such as temperature, dissolved oxygen levels, and nutrient concentrations.

In this study, we conducted a large-scale microbiome investigation of GIFT strains cultured under Malaysian climatic conditions in two distinct rearing systems: tank systems and earthen pond water, characterizing and comparing their roles on microbial communities’ structure, diversity, and variance using different biological samples. Our sampling strategy encompassed water samples from the aquatic environment where the fish were raised and all major mucosal organs, including the gut, gills, and skin. Our results reveal significant differences in microbial community structure between tank and pond water samples, reflecting each system’s contrasting environmental conditions and management practices. Furthermore, we observed distinct microbial signatures in different organ samples, indicating the presence of specialized microbial communities associated with specific anatomical niches. These findings highlight the importance of considering both host-associated and environmental factors in shaping the microbiome of aquaculture-reared fish and provide valuable insights into potential functional roles and ecological interactions within the tilapia microbiome.

## Materials and Methods

### Study Population and Sites

The Nile tilapia (*Oreochromis niloticus*) population used in this study was from a selective breeding program established in Malaysia and managed by WorldFish since 2000. The population originated from the GIFT strain selected initially for improved growth rate. To retain pedigree information, each individual was tagged with a Passive Integrated Transponder (PIT) tag at an average weight of 5 to 7g. All fish were humanely euthanized, and their body weight, standard length, total length, depth, width, sex, and PIT tag were recorded (***Figure 1A***). The fish originated from earthen ponds at Jitra Aquaculture Research Center, Kedah state (***Figure 1B***), and in Batu Maung, Penang WorldFish headquarters site in a non-recirculating tank-based system (***Figure 1C***). All fish were assessed for the presence of ectoparasite as part of routine health assessments. As described in the quick wet mount sampling guide for ectoparasites, https://hdl.handle.net/20.500.12348/4837. They were also processed for ectoparasite assessment and microbiome sampling (***Figure 1D***).

**Figure 1.**
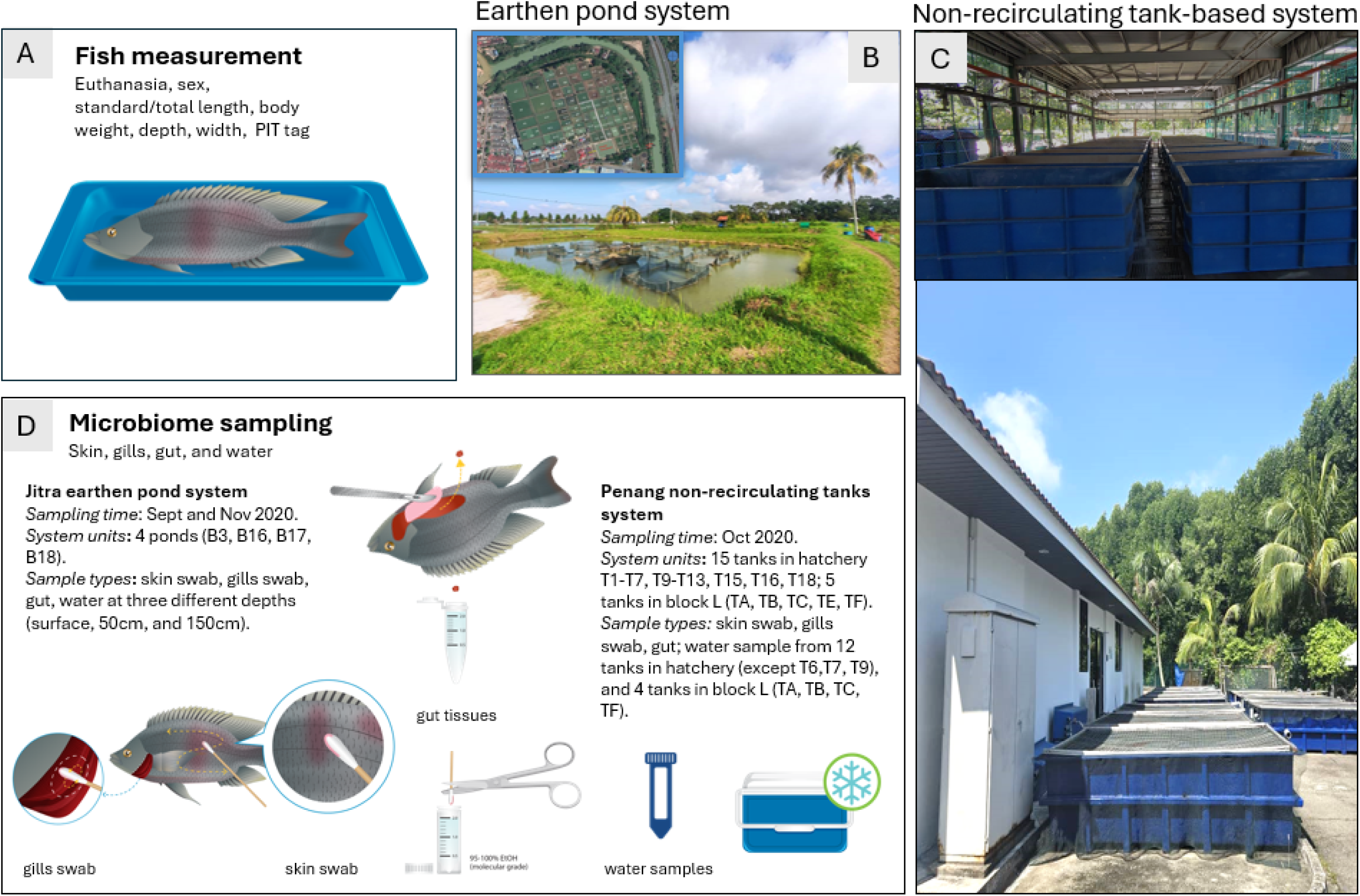
Experimental design used in this study. **A.** All fish were euthanized by an overdose of clove oil, and their length, weight, sex, and PIT tag were measured; **B.** A typical earthen pond system with hapas is at the Jitra Aquaculture Extension Center in Kedah State (inset of a satellite picture with pond layout); **C.** Typical non-recirculating tank-based system used at WorldFish headquarters site located in Batu Maung in Penang state (upper picture: tanks in hatchery covered by roof; lower picture: tanks in open air); **D.** Detailed microbiome sampling information (i.e., time, number of system units sampled, sample type) for tilapia skin, gills, gut and their water samples collected at both Jitra and Penang sites.

### Microbiome Sample Collection

Skin mucus, gills mucus, and gut samples were collected individually from all the tilapia used in this study (***Figure 1D***). Animals from Jitra were collected from four ponds, twice in September 2020 and in November 2020. Animals from Penang were collected from 20 tanks once in October 2020 (***Table 1*** and ***Supplemental Table 2***). Briefly, for skin samples, 2 sterile cotton swabs (1 swab/body side) were collected and placed in 1.5 mL screw cap tubes filled with 95% molecular grade ethanol. For the gill samples, we followed a similar procedure in a separate tube, where 2 sterile cotton swabs were used to collect the mucus from between the gill racks and filaments of at least 3 gill racks per swab for each fish (Figure 1E). For the gut, small pieces of fore-mid-hind-gut were dissected out and placed in a separate tube filled with ethanol. Water samples were collected only on two occasions, once at Jitra (September 2020) and once in Penang (October 2020) (***Table 1*** and ***Supplemental Table 2***). At Jitra, water samples were collected from four ponds at three different depths (surface, 50 cm, and 150 cm) at each pond’s corners. Each sample was collected in 50 mL falcon tubes (x 4 for each depth), kept on ice, and sent immediately to the lab for processing as described in the quick microbiome sampling guide. https://hdl.handle.net/20.500.12348/4838

**Table 1.**
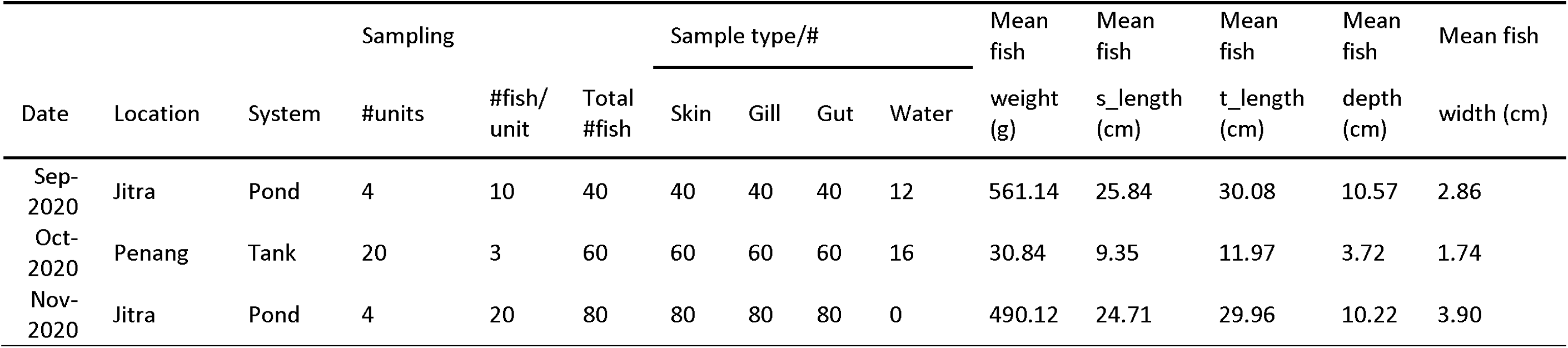
Details of sample collections.

| Date | Location | System | Sampling |  |  | Sample type/# |  |  |  | Mean fish | Mean fish | Mean fish | Mean fish | Mean fish |
| --- | --- | --- | --- | --- | --- | --- | --- | --- | --- | --- | --- | --- | --- | --- |
|  |  |  | #units | #fish/unit | Total #fish | Skin | Gill | Gut | Water | weight (g) | s_length (cm) | t_length (cm) | depth (cm) | width (cm) |
| Sep-2020 | Jitra | Pond | 4 | 10 | 40 | 40 | 40 | 40 | 12 | 561.14 | 25.84 | 30.08 | 10.57 | 2.86 |
| Oct-2020 | Penang | Tank | 20 | 3 | 60 | 60 | 60 | 60 | 16 | 30.84 | 9.35 | 11.97 | 3.72 | 1.74 |
| Nov-2020 | Jitra | Pond | 4 | 20 | 80 | 80 | 80 | 80 | 0 | 490.12 | 24.71 | 29.96 | 10.22 | 3.90 |

### DNA Extraction

Gut samples preserved in ethanol underwent centrifugation at 10,000 x g for 5 minutes, followed by the removal of the ethanol supernatant. Subsequently, 0.1 mm acid-washed silica beads were introduced into the tube containing the dissected gut sample, and homogenization was performed using a Taco Prep Bead Beater (GeneReach, Taiwan). The resulting homogenate underwent a second centrifugation at 10,000 x g for 10 minutes, with the supernatant utilized for automated nucleic acid extraction using the Taco™ Nucleic Acid Automatic Extraction System (GeneReach, Taiwan) following the manufacturer’s instructions. For skin and gill samples collected on cotton swabs and preserved in ethanol, an initial centrifugation at 10,000 x g for 5 minutes was conducted to collect any unbound or dislodged biological matters. Following the removal of the ethanol supernatant, the tubes were heated at 50°C for 30 minutes to eliminate residual ethanol on the cotton swabs. Subsequently, 200 µL of lysis buffer (1% Triton-X 100, 50 mM Tris-HCL pH 8, 5 mM EDTA, and 150 mM NaCl) was added to each swab, and the mixture was incubated at 95°C for 10 minutes (Sung et al., 2003). Then, the lysate was combined with 600 µL of chloroform and centrifuged at 10,000 x g for 5 minutes to facilitate phase separation. The upper aqueous layer was transferred to a new tube containing 1x volume Solid-phase reversible immobilization (SPRI) beads and 0.5x volume isopropanol (Oberacker et al., 2019). After incubation at room temperature for 15 minutes, the bead-bound DNA was separated on a magnetic rack, washed once with 70% ethanol, and eluted with 50 µL TE buffer. A similar methodology was applied to water samples. Briefly, 50 mL tubes containing pond or tank water were centrifuged at 4,000 rpm for 15 minutes. After removing the supernatant, the pellet was resuspended in a lysis buffer, followed by the same boiling lysis-based DNA extraction protocol described above.

### Metabarcoding: 16S V4 rRNA Amplicon Sequencing

The amplification of the 16S rRNA V4 hypervariable region employed a slightly modified partial Illumina-adapter containing forward 515F primer (underlined), 5’-TCGTCGGCAGCGTCAGATGTGTATAAGAGACAG**<u>CAGCMGCCGCGGT</u>**-3’, and 806r reverse primer, 5’-GTCTCGTGGGCTCGGAGATGTGTATAAGAGACAG**<u>GGACTACNVGGGTWTCTAAT</u>**-3’, to reduce co-amplification of host mitochondrial 12S rRNA. The PCR reaction utilized NEB OneTaq 2X Mastermix (NEB, Ipswich, MA) and followed a profile comprising an initial denaturation at 94°C for 2 minutes, followed by 30 cycles of 95°C for 15s, 47°C for 30s, and 68°C for 30s (Walters et al., 2016). The initial PCR product underwent purification using 0.8x volume SPRI beads. The cleaned PCR product was used as the template for the subsequent index PCR reaction to incorporate dual-index Nextera barcode and the remaining Illumina adapter sequence. The resulting index PCR products were pooled and separated on a 2% agarose gel. PCR band corresponding to the microbial amplicon (∼420 bp) was excised and gel-extracted using the WizBio gel purification kit (Wizbio, Korea). Library quantification of the pooled library was performed using the Denovix high-sensitivity fluorescence quantification kit (Denovix, Delaware, USA). Subsequently, the library was sequenced on an ISeq100 (Illumina, San Diego, CA) with a run configuration of 1 x 300 bp, following the recommended protocol for 16S V4 sequencing on this system (https://help.ezbiocloud.net/16s-mtp-protocol-for-illumina-iseq-100/).

### Data Analysis

Raw single-end reads were trimmed with cutadapt to remove the forward primer sequence followed by truncation of the primer-trimmed reads to a uniform length of 250 bp. Then, the reads were denoised using dada2 v1.22 (Callahan et al., 2016) as implemented within the QIIME2 pipeline (Bolyen et al., 2019). Taxonomic classification of the generated ASV used sklearn v0.22.1 (Bokulich et al., 2018) that has been trained on the recently published GreenGenes2 database (McDonald et al., 2023). The generated amplicon sequence variants (ASVs) count table and taxonomy table were subsequently exported from QIIME2 and formatted manually so that the table format is compatible with analysis using MicrobiomeAnalyst (Chong et al., 2020). During data preprocessing within the MicrobiomeAnalyst, no ASV was filtered based on prevalence and variance. All ASVs were retained for analysis to improve sensitivity, and a total sum scaling normalization was performed. Subsequent calculation and visualization of microbial relative abundance, beta and alpha-diversity calculation, and LefSE analysis based on the normalized data were performed on the MicrobiomeAnalyst web server. For the assessment of both alpha and beta diversity, statistical significance between the different groups was evaluated using permutation-based ANOVA (PerMANOVA) with 999 permutations. Linear discriminant analysis Effect Size (LefSE) analysis employed a corrected p-value cutoff of 0.05 and LDA score cutoff of 5 to identify biomarkers significantly correlated with the categories of interest.

## Associations between Descriptive Variables and Microbiomes

To determine whether the samples from various origins (sex, pond, site, tissues) exhibited markedly different microbiome profiles, we used Principal Coordinates Analysis (PCoA). PCoA is a multivariate statistical technique used to visualize similarities or differences in the microbiome abundance data by projecting distance matrices into a lower-dimensional space. In our case, the Jensen Shannon Divergence distance method was used. However, PCoA does not allow the addition of illustrative quantitative variables, i.e., all quantitative variables are necessarily used to construct the analysis. Besides, PCoA, due to its construction based on a distance matrix, it cannot take into account potential qualitative variables. However, variables such as fish sex, body weight, or age might partly explain their microbiome and parasite profiles. Thus, before constructing the PCoA, we used a PCA (R package FactoMineR v. 2.11) to assess the influence of these variables. Similarly to the PCoA, microbiome and parasite abundance data were used to construct the dimensions of the PCA. However, fish body weight and age were still projected as illustrative quantitative variables on those dimensions. Moreover, once the individuals were projected on the dimensions, the barycentres of each combination of qualitative variables (location, sampled tissue, sampling date, sex) were projected.

## Results

### Tilapia Microbiome is Shaped by Environmental Origins and Isolation Sources

A total of 568 samples were sequenced and analyzed in this study (***Table 1***). Following initial data processing, these samples underwent denoising procedures, resulting in the identification of 2,307 or amplicon sequence variants (ASVs). These ASVs represented a diverse taxonomic array, spanning 39 phyla, 86 classes, 180 orders, 299 families, 501 genera, and 399 species (***Supplemental Data 1***). A total of 4,305,794 reads were generated, with an average count per sample of 7,580, maximum of 37,796 counts per sample, and a minimum of 25 counts per sample. A total of 11 experimental factors with replicates were included (9 categoricals: sample type, location, pond, batch, loc_organ, loc_organ_pond, loc_org_month, loc_organ_month_pond, loc_month_pond; and 2 continuous, dactylogyrus and trichodina) (***Supplemental Data 2***).

When examining the Earthen pond water environment, we found that the alpha diversity of the rearing water was the highest, closely followed by the fish gill samples. In contrast, the fish gut and skin samples exhibited relatively similar average alpha diversity metrics (***Figure 2A***). Conversely, within the context of the tank water environment, the fish gill samples displayed the highest alpha diversity, followed by the tank water itself. In contrast, the gut samples exhibited the lowest alpha diversity. When categorizing based on isolation sources, including pond water, tank water, gill, gut, and skin (***Figure 2A***), both Mann-Whitney and ANOVA tests indicated significant differences in alpha-diversity values among the groups, with p-values of 2.9374e-58 and 1.2783e-94, respectively.

**Figure 2.**
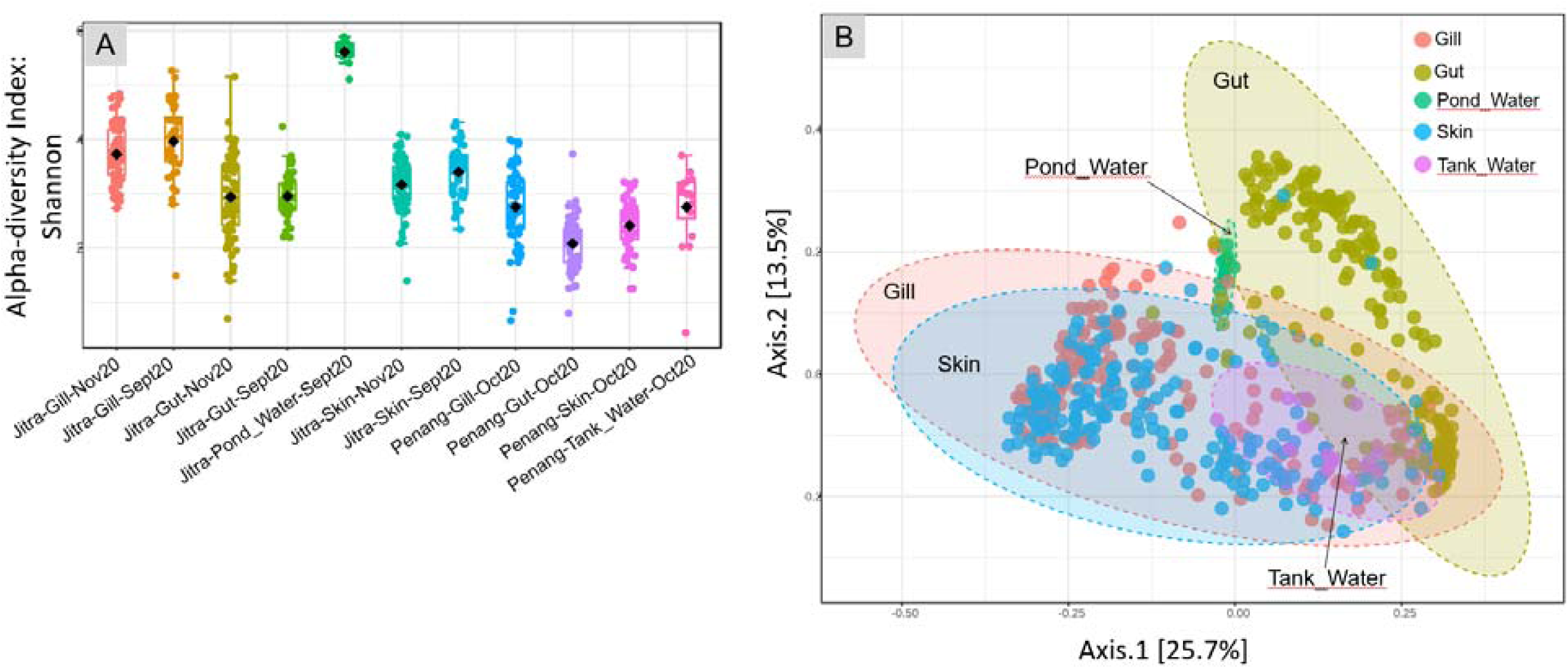
Analysis overview. **A.** Alpha diversity index: Shannon for each main group (sample type) at both Jitra and Penang locations and different time points. Statistical methods: [Mann-Whitney/Kruskal-Wallis]: p-value: 2.9374e-58; [Kruskal-Wallis] statistic: 298.69; [T-test/ANOVA]: p-value: 1.2783e-94; [ANOVA] F-value: 73.033; **B.** PCoA plot, beta-diversity that shows every sample from Jitra and Penang combined, establishing similarity between the gills, skin and water microbiomes, with gut clustering separately on the plot [ANOSIM, R: 0.3766, p-value < 0.001].

**Figure 3.**
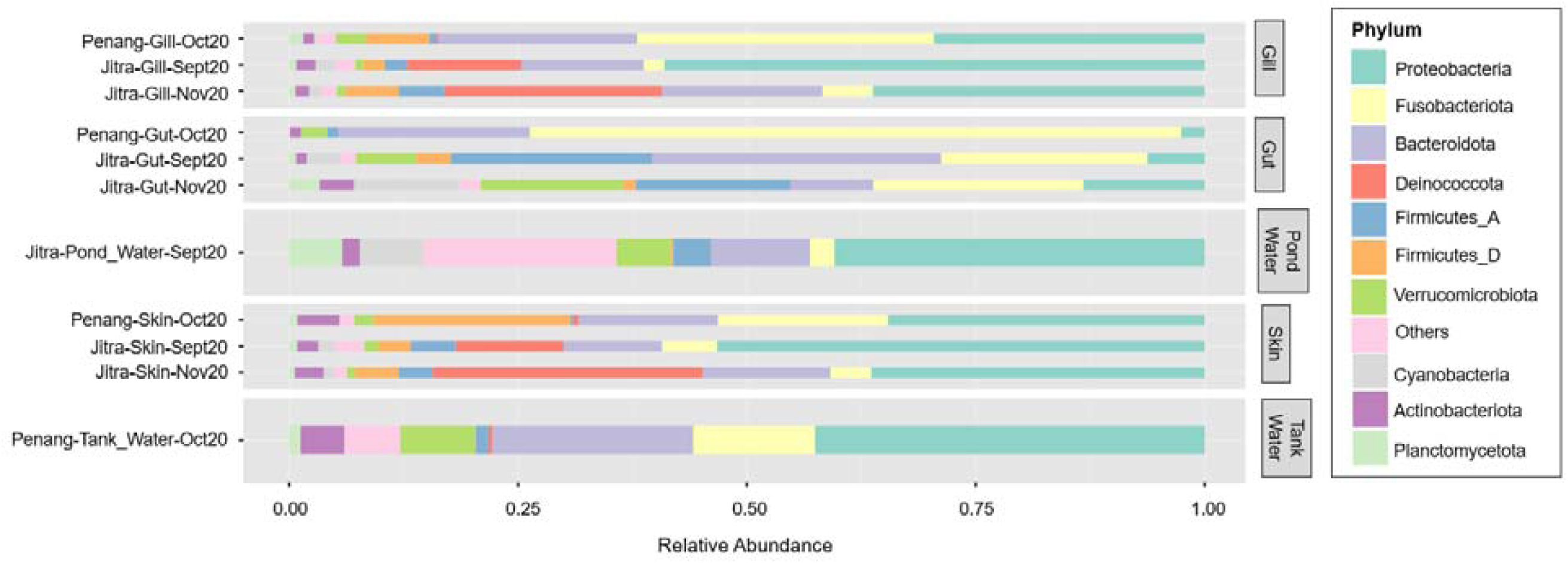
Average relative abundance of microbial phyla. Samples groups are shown along the y-axis, while phylum contributions are displayed as horizontally stacked bars, with each color representing a different phylum. The width of each segment reflects the relative abundance of that phylum within th microbial community.

A Principal Coordinates Analysis (PCoA) plot utilizing the Jensen Shannon Divergence distance method showed a clear separation of gut samples from the remaining samples (***Figure 2B***). Specifically, the fish gut samples occupied a distinct space from the lower right quadrant to the upper middle region of the plot. The separation among isolation sources was significant, as indicated by an ANOSIM value of R=0.3766 with a p-value <0.001. In contrast, samples derived from water, skin, and gill sources coalesced into a relatively expansive cluster, horizontally spanning along Axis 1, with noticeable overlap among these categories. Further stratification of the samples based on their environmental origins, specifically distinguishing between earthen ponds and fish tanks, revealed additional clustering patterns (***Figure 5***). Notably, a distinctive clustering pattern emerged within the gut samples, with fish gut samples from earthen pond water forming a discernible cluster separate from gut samples originating from fish reared in tank water (***Figure 4C***). This separation based on the rearing environment also extended to gill and skin samples. In these instances, gill and skin samples from fish reared in tank water clustered predominantly along the middle-lower right horizontal axis, while earthen pond gill and skin samples from fish reared in earthen ponds exhibited a more scattered distribution, notably concentrated in the lower left quadrant of the analysis. The separation among isolation sources was deemed significant, as indicated by an ANOSIM value of R=0.67861 with a p-value of less than 0.001.

**Figure 4.**
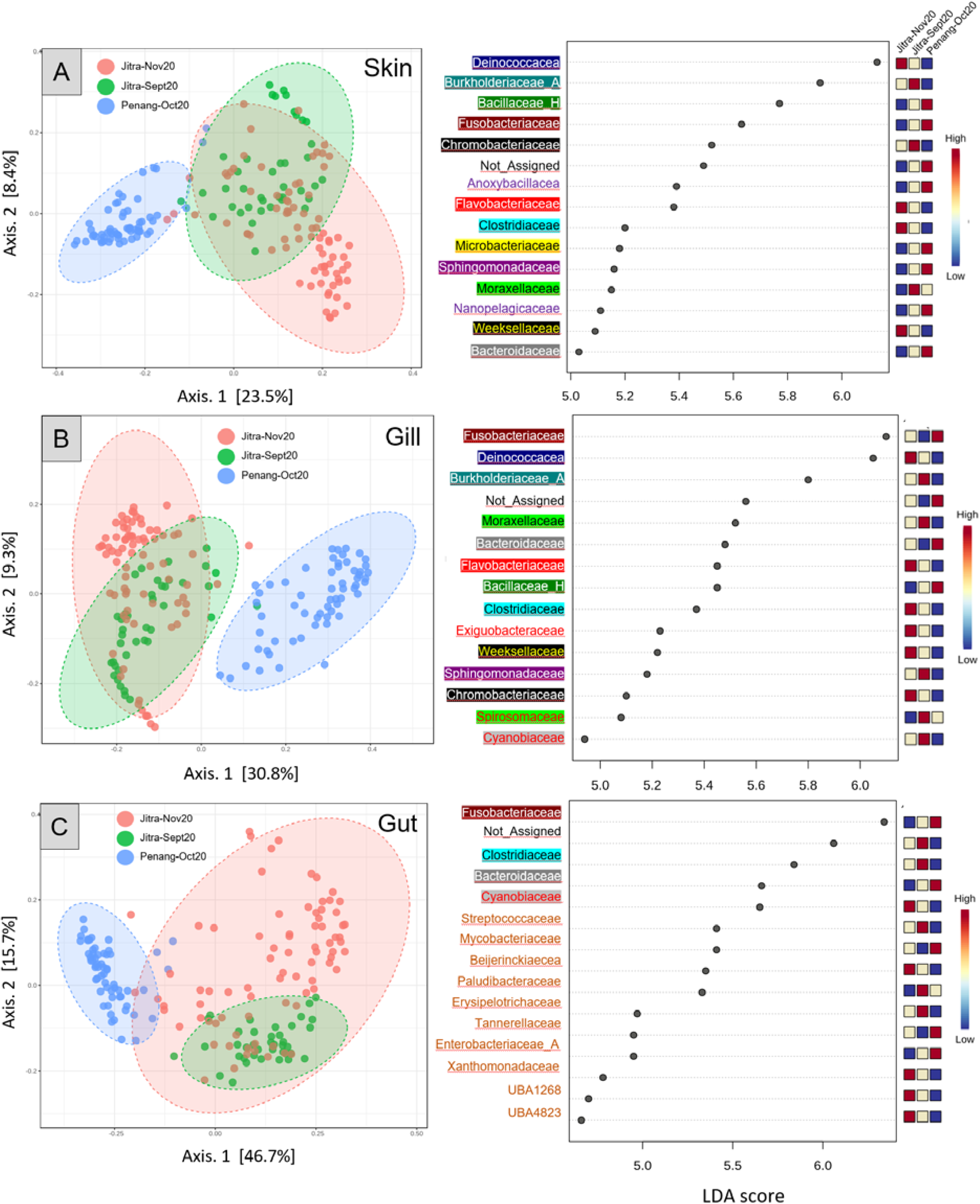
PcOA plots and LEfSe (Linear discriminant analysis Effect Size) for Jitra versus Penang skin (A), gill (B), and gut (C). Distance method: Jensen_Shannon_Divergence; Taxonomic Level: Feature level for PcOA and family level for LEfSe; *Statistical method: [ANOSIM]; Skin ANOSIM, R: 0.60735, p-value < 0.001; Gill ANOSIM, R: 0.6524, p-value < 0.001; Gut ANOSIM, R: 0.50382, p-value < 0.001.* Taxa in the LEfSe graphs were colored and highlighted based on family groups to facilitate the visualization of shared and unique taxa across different sample types.

**Figure 5.**
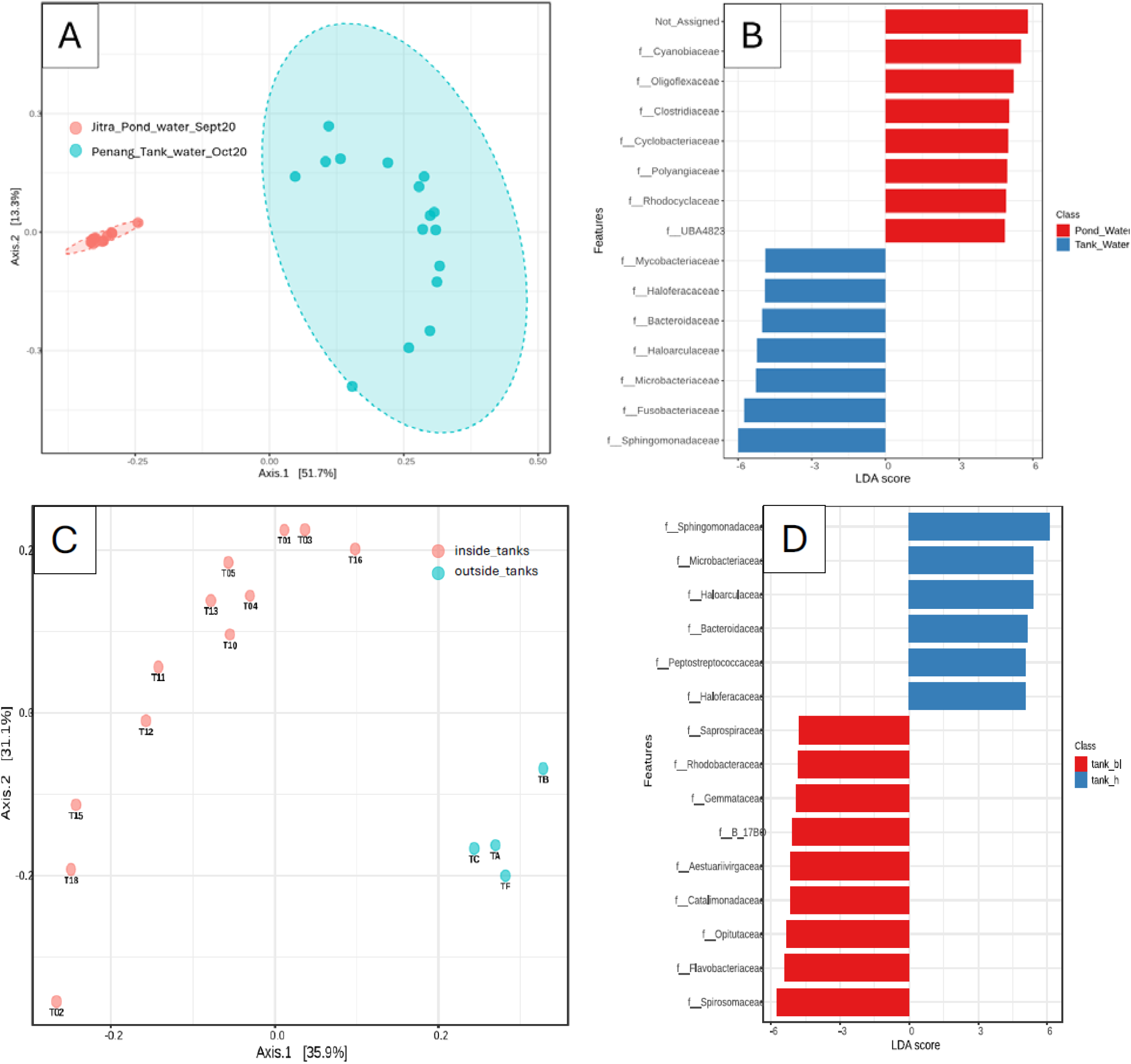
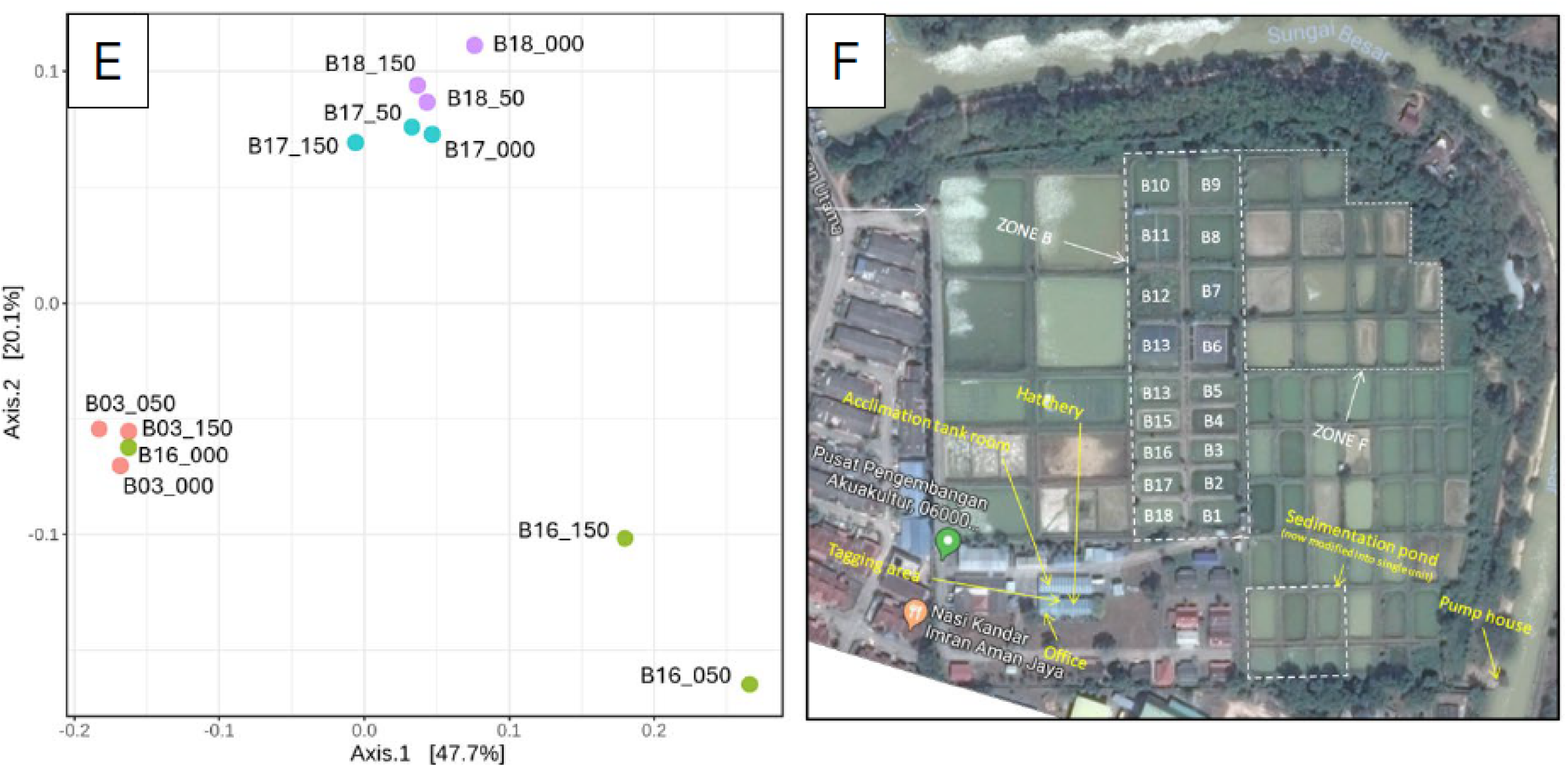
Differentiation of microbial composition in tank and pond water samples. (A) Principal Coordinate Analysis (PCoA) of microbial communities in tank and pond water samples *[ANOSIM]: 0.96, p-value < 0.001*; (B) LEfSe analysis identifying biomarkers distinguishing microbial communities between tank and pond environments; (C) PCoA of tank water samples, annotated by indoor vs. outdoor systems *[ANOSIM]: 0.767, p-value < 0.001;* (D) LEfSe analysis identifying biomarkers specific to indoor and outdoor tank systems; (E) PCoA of pond water samples annotated by pond and sampling depth *[ANOSIM]: 0.65, p-value < 0.001*; (F) Google Mapview of the earthen pond.

In the PCA, any given combination (location, sampled tissue, sampling date) was projected separately for males and females, but it always resulted in two close barycentres. (***Supplemental Data 3***). Thus, we concluded that the sex had a negligible influence on the fish microbiome and parasite profiles. Once the variables projected on the dimensions, the cos² value of the fish age and body weight illustrative variables were extracted from the PCA. It appeared that fish age cos² ranged between 2.25*10^-6^ and 0.177 on the first five dimensions, while fish body weight cos² ranged between 5.92*10^-8^ and 0.115 on the first five dimensions. These very low (< 0.2) values of cos² indicate a poor projection of fish age and body weight variables on the first five dimensions. Thus, it was concluded these two variables have also a negligible influence on the fish microbiome and parasite profiles.

### Sample Clustering at a Glance: Insights from Phyla-Level Composition

The microbial composition at the phyla level reveals clear distinctions among sample groups (***Figure 3***). Specifically, tilapia gut samples demonstrate the highest average relative abundance of reads classified under Fusobacteria. Notably, Fusobacteriota makes up a greater percentage (71.14%) of reads in gut samples from fish reared in tank water (Penang) compared to those from Earthen ponds (Jitra) (22%). Unlike Fusobacteriota, which remains relatively consistent throughout both sampling periods (September and November 2022) in Earthen pond samples, the phylum Bacteroidota experiences a substantial decrease from an average of 31.58% in September gut samples to 8.98% after two months (***Figure 3***). Furthermore, the gut samples of tilapia reared in tank water exhibit a notable absence of prominent relative abundance of reads assigned to Firmicutes_A and Cyanobacteria compared to the Earthen pond samples. While reads assigned to Firmicutes_A remain relatively consistent over the two-month period in Earthen pond gut samples, there is a substantial increase in Cyanobacteria in Jitra from September to November (3.63% to 11.49%), whereas it is nearly absent in Penang (0.03%).

Both skin and gill samples exhibit relatively similar phyla composition, reflecting the overlap observed in the PCoA plot (***Figure 2B***). There is a striking disparity in Fusobacteriota levels between the two locations (***Figure 3***). In Jitra, Fusobacteriota constitutes only 2.25% and 5.50% of the gill microbiome in September and November, respectively, whereas it represents the largest portion in Penang at 32.45%. This trend is mirrored in the skin microbiome, with an average of 18.60% of reads classified as Fusobacteriota in Penang skin samples, compared to only 6.01% and 4.44% in Jitra for September and November, respectively. On the contrary, there was a noticeable uptick in the percentage of reads classified under the phylum Deinococcota in Jitra from September (11.77% skin, 12.47% gill) to November (29.49% skin, 23.80% gill). In contrast, Deinococcota is significantly less represented by sequencing reads in Penang skin and gill samples, with only 0.57% and 0.23%, respectively.

Remarkably, Jitra’s Earthen Pond water samples exhibit a strikingly diverse and evenly distributed composition of phyla, a feature distinct from other samples. Notably, there is a substantial abundance of reads classified as Cyanobacteria and Planctomycetota in the Earthen pond water samples (7.00% and 5.76%, respectively), contrasting sharply with the Penang tank water samples (0.02% and 1.23%, respectively).

### Significant Variation in Host Microbiota between Culture Conditions and their Key Drivers

A more apparent pattern emerges after narrowing the microbiome analysis to focus on comparing samples from the same tissue source (e.g., skin, gut, gill). Analysis of the subsampled dataset using PCoA plots revealed significant differentiation among sample groups, highlighting the impact of the rearing environment on the host microbiome. Across all PCoA plots, ANOSIM values exceeded 0.5, with p-values less than 0.001, indicating strong statistical significance (Skin: R = 0.60735, p < 0.001; Gill: R = 0.6524, p < 0.001; Gut: R = 0.50382, p < 0.001; Water: R = 0.96102, p < 0.001). Although the distinction between Jitra’s September and November samples is less apparent, particularly with strong overlap observed for the skin and gill samples, a more pronounced difference is evident in the gut samples. Notably, over the two-month period, the fish gut microbiota from the Jitra undergoes significant changes, as reflected by a shift from the middle bottom part of the PCoA plot (September 2022 samples) to a more widely dispersed distribution in the upper right quadrant (November 2022 samples). Based on LEfSe analyses, the family Deinococcocea emerges with one of the highest Linear Discriminant Analysis (LDA) scores, indicating a significantly elevated representation of reads in the skin and gill samples of fish reared in earthen ponds (Figure 4A and B right). Conversely, the family Bacillacea_H, with an LDA score nearly on par with that of Deinococcacea, is notably more represented in the skin samples of fish raised in tank water (***Figure 4A right***). Moreover, among the skin and gill samples, there is a noticeable increase in the representation of reads from the family Fusobactericeae in all three types of fish microbiome from the Penang tank water system (***Figure 4A, B and C right***).

### Comparative Analysis of Microbial Communities in Tank and Pond Environment

Principal Coordinates Analysis (PCoA) of water samples obtained from tanks and ponds revealed significant and distinct clustering patterns (ANOSIM: R = 0.96102, p-value < 0.001), indicative of microbial community variation between the two aquatic environments (***Figure 5A***). Notably, the dispersion patterns observed in the PCoA plot differed markedly between tank and pond water samples. While pond water samples formed a remarkably tight cluster in the middle-left quadrant, presumably indicating a homogeneous microbial community within ponds, tank water samples exhibited a notable dispersion, occupying both the upper and lower right quadrants. Subsequent LEfSe analysis unveiled notable differences in the abundance of microbial families between pond and tank water samples (***Figure 5B***). Specifically, Cyanobiaceae exhibited the most pronounced enrichment in pond water, supported by a logarithmic Linear Discriminant Analysis (LDA) score approaching 6, while Oligoflexacea also demonstrated significant enrichment. Conversely, Sphingomonadaceae and Fusobacteriaceae were significantly enriched in tank water samples (***Figure 5B***). By focusing the analysis on solely microbial composition of tank water samples, a distinct clustering pattern emerged between water samples collected from indoor and outdoor tanks (***Figure 5C***). Notably, LefSE analysis showed that Sphingomonadaceae exhibited enrichment in samples from outside tanks, whereas Spiromaceae displayed higher abundance in inside tanks (***Figure 5D***). Interestingly, despite forming a very tight cluster among the Pond water samples (***Figure 5A***), subsequent PCOA analysis refined to only the pond water samples revealed discrete clusters by pond origin (***Figure 5E, and F***). This observation underscores the importance of targeted analyses to uncover subtle variations within seemingly homogeneous sample groups. In addition, we observed clear separations among different pond water samples but not so much when comparing the depth, suggesting that the microbiome can be a relatively “stable” indicator of pond water despite different depths.

### Potential Pathogen Load Assessment in Tilapia Amid Limited Taxonomic Resolution

Given the highly conserved nature of the 16S V4 hypervariable region, we could not identify ASVs that were classified as potential fish pathogen species, such as *Aeromonas veronii, Edwardsiella tarda, Flavobacterium columnaris,* and *S. agalactiae*. Consequently, this poses challenges for accurately analyzing pathogen load based on 16S V4 amplicon data. However, through BLAST searches of the selected ASVs from potentially pathogenic genera against the NCBI 16S rRNA refseq targeted loci database, we identified ASV15 (*Aeromonas* spp.) and ASV80 (*Edwardsiella* spp.) as potential indicators of pathogen load, as they displayed 100% nucleotide identity to multiple pathogenic species within their respective genera. For instance, ASV15 exhibited numerous identical matches to *A. media, A. jandaei, A. veronii, A. ichthiosmia* and *A. caviae*, all previously documented as opportunistic fish pathogens. Similarly, BLAST searches of ASV80 revealed identical matches to *E. tarda*. However, it is essential to note that such assumptions cannot be generalized to other genera, such as *Flavobacterium* and *Streptococcus*, as some of their members are non-pathogenic environmental or probiotic strains. The skin samples, particularly from apparently healthy Jitra fish, exhibit a higher relative abundance (∼4%) of ASV15. This is followed by gill samples, with comparatively lower relative abundances observed in gut samples. A similar trend was observed in the Penang samples albeit with even lower relative abundance across all sample types (***Figure 6***).

**Figure 6.**
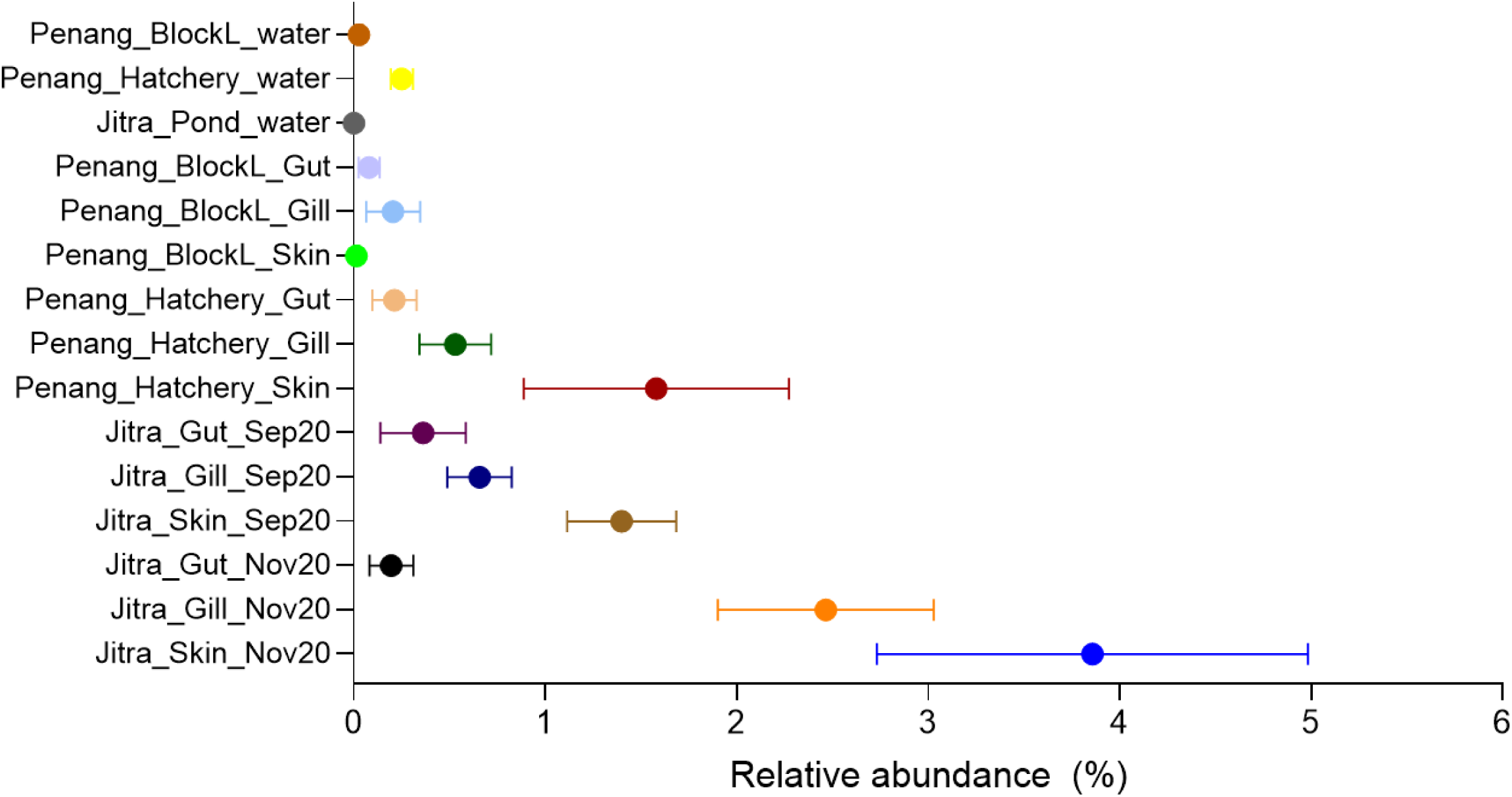
Relative abundance of ASV15 classified as *Aeromonas* in different Tilapia organs and aquaculture environments.

## Discussions

We present a comprehensive analysis of the microbiome associated with GIFT Tilapia raised in two aquaculture settings, encompassing three primary anatomical sites: gills, gut, and skin, in addition to the microbiome of their rearing water. This study provides valuable insights into the differentiation and overlap of microbiomes among fish and their immediate growth environment. Our findings indicate that pond water microbiomes consistently exhibit the highest alpha diversity, suggesting a rich diversity of bacterial taxa within this environment. This phenomenon can be attributed partly to the prevalent use of earthen ponds for tilapia cultivation in Malaysia, which are favored for their cost-effectiveness compared to cement tanks. Moreover, the sourcing of pond water from rivers introduces a diverse array of natural microbes (Tang et al., 2020), complemented by additional microbial contributions from the soil due to continuous water-soil interaction at pond bottoms, known reservoirs of microbial diversity (Colette et al., 2023; Liu et al., 2020; Wolińska et al., 2022). This heightened diversity may facilitate efficient nutrient cycling, enhancing waste conversion within the pond ecosystem. Furthermore, the daily exposure of pond water to long hours of sunlight that is naturally prevalent in this country results in a notable enrichment of cyanobacteria (Panwar et al., 2022; Zhou et al., 2014) and other aquatic plant species that serve as an additional nutritional source for fish or and other pond organisms, thereby diversifying nutrient availability beyond traditional fish feed. In contrast, tank water microbiomes exhibit significantly lower alpha diversity, likely attributed to factors such as pre-chlorination of town water sources before use and weekly water exchanges, which disrupt microbial communities before they can stabilize, thereby limiting diversity (Bertelli et al., 2018; Zhang et al., 2022; Zhou et al., 2021). While this may mitigate the spread of pathogens, it also reduces overall microbiome diversity. This shows there may be trade-offs in terms of health and growth in these different aquaculture systems.

Despite continuous exposure and interaction with the water microbiome, the microbiomes of the tilapia tissue types sampled exhibited distinct gut microbial compositions, indicative of host-driven microbial selection. The gut microbiome consistently demonstrates lower diversity than other organs, a well-established observation in the scientific literature (Abdelhafiz, 2022; Parata et al., 2020; Wu et al., 2021). However, it is intriguing to note that the gut microbiome diversity in Jitra samples is generally higher than that of tank-reared counterparts, likely influenced by dietary factors (do Vale Pereira et al., 2024; Escalas et al., 2022; Soh et al., 2024). While tank-reared fish adhere to controlled feeding regimes, fish reared in earthen ponds, take water by limited drinking and benefit from a diet supplemented by materials available within the pond environment, such as algae and aquatic plant species. This is supported by the increased abundance of Firmicutes_A in Jitra fish guts, reported for their role in plant carbohydrate degradation (Berry, 2016; do Vale Pereira et al., 2024; Flint et al., 2012). Nevertheless, considering the age difference among fish from both sites, we cannot discount the possibility that microbiome differences may also be influenced by age (Zhao et al., 2020). As expected, a high number of reads were assigned to Fusobacteria, particularly the genus *Cetobacterium,* in the fish gut from both systems. *Cetobacterium*, a dominant genus in freshwater fish gut microbiota, plays key roles in gut fermentation processes and vitamin B12 synthesis for the host (Colorado Gómez et al., 2023; Ofek et al., 2021; Qi et al., 2023; Yajima et al., 2023). In addition, Fusobacteriota is recognized for its capacity to produce butyrate, a short-chain fatty acid well-known for its many benefits to the host. Butyrate serves as an anti-inflammatory agent, promotes mucus secretion, and provides energy to host cells (Larsen et al., 2014; von Engelhardt et al., 1998). Moreover, butyric acid has been reported to inhibit freshwater fish pathogens such as *Aeromonas* sp., *Flavobacterium* sp., *Yersinia* sp., and *Vibrio* sp.) (Larsen et al., 2014). That said, the increased abundance of Fusobacteria in gill and water samples from tank-reared fish is unlikely to be biologically driven but rather reflects tank operation, such as the circulation of fecal material within the tank environment driven by aeration in the absence of a physical filtration system. Gill structures, crucial for respiratory water passage, occasionally trap and sample these fecal materials, contributing to the observed microbial composition.

The prevalence of Deinoccota in the skin and gill microbiome of fish in Jitra ponds over two distinct time points spaced 60 days apart is intriguing, given that Deinoccota is not commonly reported to be present in high abundance on the skin or gill microbiome of Tilapia (Berggren et al., 2022; Debnath et al., 2023; McMurtrie et al., 2022). Members of the phylum Deinoccocata, particularly within the family Deinoccocacea, are known for their exceptional resistance to extreme radiation, including UV radiation (Liu et al., 2023; Zhang et al., 2007), and are commonly found in soil microbiomes (Lang et al., 2023; Luo et al., 2021; Rainey et al., 2005). Therefore, the direct interaction of pond water with underlying soil in Jitra may provide a continuous reservoir of Deinoccota, facilitating their colonization of the fish skin microbiome. The presence of Deinoccota on the fish’s skin may serve two functional roles. Firstly, their resistance to UV radiation suggests a potential protective mechanism for the fish against sunlight exposure (Farci et al., 2016), particularly relevant in environments with continuous sunlight exposure, such as Jitra ponds. Secondly, Deinoccota may contribute to the establishment and maintenance of a stable microbiome on the fish skin, potentially through niche occupation and competitive interactions with other microorganisms, thereby aiding in the prevention of pathogen colonization and promoting a balanced microbial community (Cabillon and Lazado, 2019; Larsen et al., 2013; Takeuchi et al., 2021). The skin and gill of fish originating from either earthen pond or tank system had similar bacterial composition as reflected by their overlapping beta-diversity profiles and relative abundance. Those similarities could be explained by the fact that these two mucosal organs are in constant contact with their external water environment. Other studies have reported similar observations in tilapia co-cultivated with grey-mullet (Elsheshtawy et al., 2021) and in many other fish species, such as Atlantic salmon (Lorgen-Ritchie et al., 2022) and yellowtail kingfish (Legrand et al., 2018).

Belonging to the Proteobacteria phylum, *Aeromonas* encompasses several species recognized as potential opportunistic pathogens of tilapia, including *A. hydrophila, jandaei,* and *veronii* (Basri et al., 2020; Dong et al., 2017). However, *Aeromonas* has also been identified in the healthy intestinal mucosa of tilapia, suggesting its potential role in maintaining and stimulating mucosal immunity (Elsheshtawy et al., 2021; Nayak, 2010; Pakingking et al., 2015; Wu et al., 2021). In our study, we observed the presence of *Aeromonas* in healthy tilapia, albeit in low relative abundance. Predominantly, reads assigned to *Aeromonas* were found in the skin tissues, followed by gill samples, and to a lesser extent in the gut of fish within earthen-pond environments, with minimal occurrences in fish from tank systems, except for isolated instances in specific tanks. In farm settings, effective management of *Aeromonas* is pivotal to maintaining their levels below a critical threshold, particularly considering various stressors like poor water quality that could disrupt the delicate balance of the microbiome, rendering the animals more susceptible to *Aeromonas* and other potential opportunistic pathogens. Traditional molecular detection techniques of *Aeromonas,* such as standard PCR, often lack quantitative data, potentially leading to false alarms for producers. Consequently, the 16S rRNA amplicon sequencing method emerges as a complementary tool, providing farmers with an enhanced understanding of microbial communities and their relative abundance, especially concerning potential opportunistic pathogens. In addition, the adoption of full-length 16S rRNA gene sequencing for fish microbiomes using the Nanopore sequencing platform will enable researchers and producers to categorize the microbial communities with species-level precision rapidly.

In conclusion, our study sheds light on the intricate dynamics of the microbiome associated with GIFT Tilapia in two different aquaculture settings. By examining microbial communities across various fish organs and water environments, we unveil significant differences in microbial diversity and structure between pond and tank systems, highlighting the influence of environmental factors such as water source and management practices. Microbiome profiling holds significant promise for enhancing routine decision-making in pond management and bolstering aquaculture sustainability. However, its full potential can only be realized if it is readily accessible to farmers in a timely manner. Therefore, streamlining sampling, library preparation, and sequencing procedures are paramount. Encouragingly, the rapid advancements in the portable nanopore sequencing platforms offer hope for transformative change in microbiome profiling within aquaculture. By facilitating swift and cost-effective analysis, these innovations have the potential to revolutionize aquaculture systems, leading to increased efficiency and productivity in the industry.

## Ethics approval and consent to participate

Data collection and sampling were performed as part of a non-profit selective breeding program run by WorldFish, where the samples were collected on euthanized fish as part of routine husbandry procedures, i.e., population health checks and stocking density adjustments. The animals from this breeding population are managed in accordance with the Guiding Principles of the Animal Care, Welfare, and Ethics Policy of WorldFish Center.

## Data availability

FastQ files for all 568 samples can be found under BioProject accession PRJNA1036009 with the following BioSamples accession from SAMN38257401 to SAMN38257968. The Sequence Read Archive (SRA) accession numbers for all 568 samples can be found in ***Supplemental Data 4*.**

## Supporting information

Supplemental Data1_ASV_Sequence

Supplemental_Data2_Microbiome_input_files

Supplemental_Data3_PCA

Supplemental_Data4_SRAs

## Acknowledgments

The authors gratefully acknowledge financial support from the CGIAR Research Program on Fish Agri-Food Systems (FISH), the CGIAR Initiative on “Aquatic Foods” led by WorldFish and “Protecting Human Health Through a One Health Approach”. This research also receive support from Norway as part of the Aquatic Animal Health Africa project “Increased Sustainability in the Aquaculture Sector in sub-Saharan Africa, through Improved Aquatic Animal Health Management” led by WorldFish and the Norwegian Veterinary Institute [grant no. RAF-19/0051]. We thank all funders who support this research through their contributions to the CGIAR Trust Fund: https://www.cgiar.org/funders/, and Norway. Funders were not involved in the conceptualization, design, data collection, analysis, decision to publish, or manuscript preparation. We thank the WorldFish breeding team and the Department of Fisheries Malaysia for their technical support throughout the project. The contents of this publication are the sole responsibility of the authors and can in no way be taken to reflect the views of the Government of Norway.

## Authors’ contributions

JDD, VMC, JAHB: Conceptualization; JDD, MM, HMG: Data curation; JDD, HMG, CR: Formal analysis; JDD, MM, LK, VMC, JAHB: Investigation; JDD, MM, HMG, CR, LK, VMC, JAHB: Methodology; JDD, MM, HMG, CR: Visualization; JDD, HMG: Writing-original draft; JDD, MM, HMG, CR, LK, DVJ, VMC, JAHB: Writing – review & editing. All authors improved the writing, read and approved the final manuscript.

## Supplemental Data Description

- Supplemental Data 1. ASV Sequences in fasta format generated in this study
- Supplemental Data 2. MicrobiomeAnalyst input files with various metadata combinations
- Supplemental Data 3. PCA, coordinates of the barycenters of the PCA, and correlation and cos² values of Fish Age and Body Weight
- Supplemental Data 4. List of SRA accession codes for each raw read

## References

1. Abdelhafiz, Y.A.A., 2022. Insights into the bacterial communities of Nile tilapia – core members and intergenerational transfer (Doctoral thesis). 106. Nord University.

2. Barría, A., Trinh, T.Q., Mahmuddin, M., Benzie, J.A.H., Chadag, V.M., Houston, R.D., 2020. Genetic parameters for resistance to Tilapia Lake Virus (TiLV) in Nile tilapia (Oreochromis niloticus). Aquaculture 522, 735126. 10.1016/j.aquaculture.2020.735126

3. Basri, L., Nor, R.M., Salleh, A., Md. Yasin, I.S., Saad, M.Z., Abd. Rahaman, N.Y., Barkham, T., Amal, M.N.A., 2020. Co-Infections of Tilapia Lake Virus, Aeromonas hydrophila and Streptococcus agalactiae in Farmed Red Hybrid Tilapia. Animals 10, 2141. 10.3390/ani10112141

4. Berggren, H., Tibblin, P., Yıldırım, Y., Broman, E., Larsson, P., Lundin, D., Forsman, A., 2022. Fish Skin Microbiomes Are Highly Variable Among Individuals and Populations but Not Within Individuals. Frontiers in Microbiology 12.

5. Berry, D., 2016. The emerging view of Firmicutes as key fibre degraders in the human gut. Environmental Microbiology 18, 2081–2083. 10.1111/1462-2920.13225

6. Bertelli, C., Courtois, S., Rosikiewicz, M., Piriou, P., Aeby, S., Robert, S., Loret, J.-F., Greub, G., 2018. Reduced Chlorine in Drinking Water Distribution Systems Impacts Bacterial Biodiversity in Biofilms. Front. Microbiol. 9. 10.3389/fmicb.2018.02520

7. Bokulich, N.A., Kaehler, B.D., Rideout, J.R., Dillon, M., Bolyen, E., Knight, R., Huttley, G.A., Gregory Caporaso, J., 2018. Optimizing taxonomic classification of marker-gene amplicon sequences with QIIME 2’s q2-feature-classifier plugin. Microbiome 6, 90. 10.1186/s40168-018-0470-z

8. Bolyen, E., Rideout, J.R., Dillon, M.R., Bokulich, N.A., Abnet, C.C., Al-Ghalith, G.A., Alexander, H., Alm, E.J., Arumugam, M., Asnicar, F., Bai, Y., Bisanz, J.E., Bittinger, K., Brejnrod, A., Brislawn, C.J., Brown, C.T., Callahan, B.J., Caraballo-Rodríguez, A.M., Chase, J., Cope, E.K., Da Silva, R., Diener, C., Dorrestein, P.C., Douglas, G.M., Durall, D.M., Duvallet, C., Edwardson, C.F., Ernst, M., Estaki, M., Fouquier, J., Gauglitz, J.M., Gibbons, S.M., Gibson, D.L., Gonzalez, A., Gorlick, K., Guo, J., Hillmann, B., Holmes, S., Holste, H., Huttenhower, C., Huttley, G.A., Janssen, S., Jarmusch, A.K., Jiang, L., Kaehler, B.D., Kang, K.B., Keefe, C.R., Keim, P., Kelley, S.T., Knights, D., Koester, I., Kosciolek, T., Kreps, J., Langille, M.G.I., Lee, J., Ley, R., Liu, Y.-X., Loftfield, E., Lozupone, C., Maher, M., Marotz, C., Martin, B.D., McDonald, D., McIver, L.J., Melnik, A.V., Metcalf, J.L., Morgan, S.C., Morton, J.T., Naimey, A.T., Navas-Molina, J.A., Nothias, L.F., Orchanian, S.B., Pearson, T., Peoples, S.L., Petras, D., Preuss, M.L., Pruesse, E., Rasmussen, L.B., Rivers, A., Robeson, M.S., Rosenthal, P., Segata, N., Shaffer, M., Shiffer, A., Sinha, R., Song, S.J., Spear, J.R., Swafford, A.D., Thompson, L.R., Torres, P.J., Trinh, P., Tripathi, A., Turnbaugh, P.J., Ul-Hasan, S., Van Der Hooft, J.J.J., Vargas, F., Vázquez-Baeza, Y., Vogtmann, E., Von Hippel, M., Walters, W., Wan, Y., Wang, M., Warren, J., Weber, K.C., Williamson, C.H.D., Willis, A.D., Xu, Z.Z., Zaneveld, J.R., Zhang, Y., Zhu, Q., Knight, R., Caporaso, J.G., 2019. Reproducible, interactive, scalable and extensible microbiome data science using QIIME 2. Nat Biotechnol 37, 852–857. 10.1038/s41587-019-0209-9

9. Cabillon, N.A.R., Lazado, C.C., 2019. Mucosal Barrier Functions of Fish under Changing Environmental Conditions. Fishes 4, 2. 10.3390/fishes4010002

10. Callahan, B.J., McMurdie, P.J., Rosen, M.J., Han, A.W., Johnson, A.J.A., Holmes, S.P., 2016. DADA2: High-resolution sample inference from Illumina amplicon data. Nat Methods 13, 581–583. 10.1038/nmeth.3869

11. Chong, J., Liu, P., Zhou, G., Xia, J., 2020. Using MicrobiomeAnalyst for comprehensive statistical, functional, and meta-analysis of microbiome data. Nat Protoc 15, 799–821. 10.1038/s41596-019-0264-1

12. Colette, M., Guentas, L., Patrona, L.D., Ansquer, D., Callac, N., 2023. Dynamic of active microbial diversity in rhizosphere sediments of halophytes used for bioremediation of earthen shrimp ponds. Environmental Microbiome 18, 58. 10.1186/s40793-023-00512-x

13. Colorado Gómez, M.A., Melo-Bolívar, J.F., Ruíz Pardo, R.Y., Rodriguez, J.A., Villamil, L.M., 2023. Unveiling the Probiotic Potential of the Anaerobic Bacterium Cetobacterium sp. nov. C33 for Enhancing Nile Tilapia (Oreochromis niloticus) Cultures. Microorganisms 11, 2922. 10.3390/microorganisms11122922

14. Debnath, S.C., McMurtrie, J., Temperton, B., Delamare-Deboutteville, J., Mohan, C.V., Tyler, C.R., 2023. Tilapia aquaculture, emerging diseases, and the roles of the skin microbiomes in health and disease. Aquacult Int. 10.1007/s10499-023-01117-4

15. do Vale Pereira, G., Teixeira, C., Couto, J., Dias, J., Rema, P., Gonçalves, A.T., 2024. Dietary Protein Quality Affects the Interplay between Gut Microbiota and Host Performance in Nile Tilapia. Animals 14, 714. 10.3390/ani14050714

16. Dong, H.T., Chaijarasphong, T., Barnes, A.C., Delamare-Deboutteville, J., Lee, P.A., Senapin, S., Mohan, C.V., Tang, K.F.J., McGladdery, S.E., Bondad-Reantaso, M.G., 2023. From the basics to emerging diagnostic technologies: What is on the horizon for tilapia disease diagnostics? Reviews in Aquaculture 15, 186–212. 10.1111/raq.12734

17. Dong, H.T., Techatanakitarnan, C., Jindakittikul, P., Thaiprayoon, A., Taengphu, S., Charoensapsri, W., Khunrae, P., Rattanarojpong, T., Senapin, S., 2017. Aeromonas jandaei and Aeromonas veronii caused disease and mortality in Nile tilapia, Oreochromis niloticus (L.). J Fish Dis 40, 1395–1403. 10.1111/jfd.12617

18. Elsheshtawy, A., Clokie, B.G.J., Albalat, A., Beveridge, A., Hamza, A., Ibrahim, A., MacKenzie, S., 2021. Characterization of External Mucosal Microbiomes of Nile Tilapia and Grey Mullet Co-cultured in Semi-Intensive Pond Systems. Frontiers in Microbiology 12.

19. Escalas, A., Auguet, J.-C., Avouac, A., Belmaker, J., Dailianis, T., Kiflawi, M., Pickholtz, R., Skouradakis, G., Villéger, S., 2022. Shift and homogenization of gut microbiome during invasion in marine fishes. Animal Microbiome 4, 37. 10.1186/s42523-022-00181-0

20. Etherington, G.J., Nash, W., Ciezarek, A., Mehta, T.K., Barria, A., Peñaloza, C., Khan, M.G.Q., Durrant, A., Forrester, N., Fraser, F., Irish, N., Kaithakottil, G.G., Lipscombe, J., Trong, T., Watkins, C., Swarbreck, D., Angiolini, E., Cnaani, A., Gharbi, K., Houston, R.D., Benzie, J. a. H., Haerty, W., 2022. Chromosome-level genome sequence of the Genetically Improved Farmed Tilapia (GIFT, Oreochromis niloticus) highlights regions of introgression with O. mossambicus. BMC Genomics 23, 832. 10.1186/s12864-022-09065-8

21. FAO, 2024. The State of World Fisheries and Aquaculture 2024. Blue Transformation in action. Rome.

22. Farci, D., Slavov, C., Tramontano, E., Piano, D., 2016. The S-layer Protein DR_2577 Binds Deinoxanthin and under Desiccation Conditions Protects against UV-Radiation in Deinococcus radiodurans. Front. Microbiol. 7. 10.3389/fmicb.2016.00155

23. Flint, H.J., Scott, K.P., Duncan, S.H., Louis, P., Forano, E., 2012. Microbial degradation of complex carbohydrates in the gut. Gut Microbes 3, 289–306. 10.4161/gmic.19897

24. Hamzah, A., Ponzoni, R.W., Nguyen, N.H., Khaw, H., Yee, H.Y., Mohd Nor, S.A., 2014. Performance of the Genetically Improved Farmed Tilapia (GIFT) strain over ten generations of selection in Malaysia.

25. Lang, Y., Ma, Y., Wang, G., Qian, X., Wang, J., 2023. Optimized Fertilization Shifted Soil Microbial Properties and Improved Vegetable Growth in Facility Soils with Obstacles. Horticulturae 9, 1303. 10.3390/horticulturae9121303

26. Larsen, A., Tao, Z., Bullard, S.A., Arias, C.R., 2013. Diversity of the skin microbiota of fishes: evidence for host species specificity. FEMS Microbiology Ecology 85, 483–494. 10.1111/1574-6941.12136

27. Larsen, A.M., Mohammed, H.H., Arias, C.R., 2014. Characterization of the gut microbiota of three commercially valuable warmwater fish species. J Appl Microbiol 116, 1396–1404. 10.1111/jam.12475

28. Legrand, T.P.R.A., Catalano, S.R., Wos-Oxley, M.L., Stephens, F., Landos, M., Bansemer, M.S., Stone, D.A.J., Qin, J.G., Oxley, A.P.A., 2018. The Inner Workings of the Outer Surface: Skin and Gill Microbiota as Indicators of Changing Gut Health in Yellowtail Kingfish. Front Microbiol 8, 2664. 10.3389/fmicb.2017.02664

29. Liu, S., Wang, F., Chen, H., Yang, Z., Ning, Y., Chang, C., Yang, D., 2023. New Insights into Radio-Resistance Mechanism Revealed by (Phospho)Proteome Analysis of Deinococcus Radiodurans after Heavy Ion Irradiation. International Journal of Molecular Sciences 24, 14817. 10.3390/ijms241914817

30. Liu, Z., Iqbal, M., Zeng, Z., Lian, Y., Zheng, A., Zhao, M., Li, Zixin, Wang, G., Li, Zhifei, Xie, J., 2020. Comparative analysis of microbial community structure in the ponds with different aquaculture model and fish by high-throughput sequencing. Microbial Pathogenesis 142, 104101. 10.1016/j.micpath.2020.104101

31. Lorgen-Ritchie, M., Clarkson, M., Chalmers, L., Taylor, J.F., Migaud, H., Martin, S.A.M., 2022. Temporal changes in skin and gill microbiomes of Atlantic salmon in a recirculating aquaculture system – Why do they matter? Aquaculture 558, 738352. 10.1016/j.aquaculture.2022.738352

32. Luo, Y., Yuan, H., Zhao, J., Qi, Y., Cao, W.-W., Liu, J.-M., Guo, W., Bao, Z.-H., 2021. Multiple factors influence bacterial community diversity and composition in soils with rare earth element and heavy metal co-contamination. Ecotoxicology and Environmental Safety 225, 112749. 10.1016/j.ecoenv.2021.112749

33. McDonald, D., Jiang, Y., Balaban, M., Cantrell, K., Zhu, Q., Gonzalez, A., Morton, J.T., Nicolaou, G., Parks, D.H., Karst, S.M., Albertsen, M., Hugenholtz, P., DeSantis, T., Song, S.J., Bartko, A., Havulinna, A.S., Jousilahti, P., Cheng, S., Inouye, M., Niiranen, T., Jain, M., Salomaa, V., Lahti, L., Mirarab, S., Knight, R., 2023. Greengenes2 unifies microbial data in a single reference tree. Nat Biotechnol 1–4. 10.1038/s41587-023-01845-1

34. McMurtrie, J., Alathari, S., Chaput, D.L., Bass, D., Ghambi, C., Nagoli, J., Delamare-Deboutteville, J., Mohan, C.V., Cable, J., Temperton, B., Tyler, C.R., 2022. Relationships between pond water and tilapia skin microbiomes in aquaculture ponds in Malawi. Aquaculture 558, 738367. 10.1016/j.aquaculture.2022.738367

35. Nayak, S.K., 2010. Role of gastrointestinal microbiota in fish. Aquaculture Research 41, 1553– 1573. 10.1111/j.1365-2109.2010.02546.x

36. Oberacker, P., Stepper, P., Bond, D.M., Höhn, S., Focken, J., Meyer, V., Schelle, L., Sugrue, V.J., Jeunen, G.-J., Moser, T., Hore, S.R., Meyenn, F. von, Hipp, K., Hore, T.A., Jurkowski, T.P., 2019. Bio-On-Magnetic-Beads (BOMB): Open platform for high-throughput nucleic acid extraction and manipulation. PLOS Biology 17, e3000107. 10.1371/journal.pbio.3000107

37. Ofek, T., Lalzar, M., Laviad-Shitrit, S., Izhaki, I., Halpern, M., 2021. Comparative Study of Intestinal Microbiota Composition of Six Edible Fish Species. Front. Microbiol. 12. 10.3389/fmicb.2021.760266

38. Pakingking, R., Palma, P., Usero, R., 2015. Quantitative and qualitative analyses of the bacterial microbiota of tilapia (Oreochromis niloticus) cultured in earthen ponds in the Philippines. World J Microbiol Biotechnol 31, 265–275. 10.1007/s11274-014-1758-1

39. Panwar, P., Williams, T.J., Allen, M.A., Cavicchioli, R., 2022. Population structure of an Antarctic aquatic cyanobacterium. Microbiome 10, 207. 10.1186/s40168-022-01404-x

40. Parata, L., Mazumder, D., Sammut, J., Egan, S., 2020. Diet type influences the gut microbiome and nutrient assimilation of Genetically Improved Farmed Tilapia (Oreochromis niloticus). PLoS One 15, e0237775. 10.1371/journal.pone.0237775

41. Ponzoni, R.W., Nguyen, N.H., Khaw, H.L., Hamzah, A., Bakar, K.R.A., Yee, H.Y., 2011. Genetic improvement of Nile tilapia (Oreochromis niloticus) with special reference to the work conducted by the WorldFish Center with the GIFT strain. Reviews in Aquaculture 3, 27–41. 10.1111/j.1753-5131.2010.01041.x

42. Qi, X., Zhang, Yong, Zhang, Yilin, Luo, F., Song, K., Wang, G., Ling, F., 2023. Vitamin B12 produced by Cetobacterium somerae improves host resistance against pathogen infection through strengthening the interactions within gut microbiota. Microbiome 11, 135. 10.1186/s40168-023-01574-2

43. Rainey, F.A., Ray, K., Ferreira, M., Gatz, B.Z., Nobre, M.F., Bagaley, D., Rash, B.A., Park, M.-J., Earl, A.M., Shank, N.C., Small, A.M., Henk, M.C., Battista, J.R., Kämpfer, P., da Costa, M.S., 2005. Extensive Diversity of Ionizing-Radiation-Resistant Bacteria Recovered from Sonoran Desert Soil and Description of Nine New Species of the Genus Deinococcus Obtained from a Single Soil Sample. Applied and Environmental Microbiology 71, 5225–5235. 10.1128/AEM.71.9.5225-5235.2005

44. Soh, M., Tay, Y.C., Lee, C.S., Low, A., Orban, L., Jaafar, Z., Seedorf, H., 2024. The intestinal digesta microbiota of tropical marine fish is largely uncultured and distinct from surrounding water microbiota. npj Biofilms Microbiomes 10, 1–15. 10.1038/s41522-024-00484-x

45. Sung, K., Khan, S.A., Nawaz, M.S., Khan, A.A., 2003. A simple and efficient Triton X-100 boiling and chloroform extraction method of RNA isolation from Gram-positive and Gram-negative bacteria. FEMS Microbiol Lett 229, 97–101. 10.1016/S0378-1097(03)00791-2

46. Tacon, A.G.J., Shumway, S.E., 2024. Critical Need to Increase Aquatic Food Production and Food Supply from Aquaculture and Capture Fisheries: Trends and Outlook. Reviews in Fisheries Science & Aquaculture 32, 389–395. 10.1080/23308249.2024.2324321

47. Takeuchi, M., Fujiwara-Nagata, E., Katayama, T., Suetake, H., 2021. Skin bacteria of rainbow trout antagonistic to the fish pathogen Flavobacterium psychrophilum. Sci Rep 11, 7518. 10.1038/s41598-021-87167-1

48. Tang, X., Xie, G., Shao, K., Hu, Y., Cai, J., Bai, C., Gong, Y., Gao, G., 2020. Contrast diversity patterns and processes of microbial community assembly in a river-lake continuum across a catchment scale in northwestern China. Environmental Microbiome 15, 10. 10.1186/s40793-020-00356-9

49. von Engelhardt, W., Bartels, J., Kirschberger, S., Meyer zu Düttingdorf, H.D., Busche, R., 1998. Role of short-chain fatty acids in the hind gut. Vet Q 20 Suppl 3, S52–59.

50. Walters, W., Hyde, E.R., Berg-Lyons, D., Ackermann, G., Humphrey, G., Parada, A., Gilbert, J.A., Jansson, J.K., Caporaso, J.G., Fuhrman, J.A., Apprill, A., Knight, R., 2016. Improved Bacterial 16S rRNA Gene (V4 and V4-5) and Fungal Internal Transcribed Spacer Marker Gene Primers for Microbial Community Surveys. mSystems 1, e00009–15. 10.1128/mSystems.00009-15

51. Wolińska, A., Ndongo, A., Grządziel, J., Gałązka, G., Marzec-Grządziel, A., Szałaj, K., Kuźniar, K., 2022. Functional and Seasonal Changes in the Structure of Microbiome Inhabiting Bottom Sediments of a Pond Intended for Ecological King Carp Farming. Biology 11. 10.3390/biology11060913

52. Wu, Z., Zhang, Q., Lin, Y., Hao, J., Wang, S., Zhang, J., Li, A., 2021. Taxonomic and Functional Characteristics of the Gill and Gastrointestinal Microbiota and Its Correlation with Intestinal Metabolites in NEW GIFT Strain of Farmed Adult Nile Tilapia (Oreochromis niloticus). Microorganisms 9, 617. 10.3390/microorganisms9030617

53. Yajima, D., Fujita, H., Hayashi, I., Shima, G., Suzuki, K., Toju, H., 2023. Core species and interactions prominent in fish-associated microbiome dynamics. Microbiome 11, 53. 10.1186/s40168-023-01498-x

54. Zhang, Yu-Qin, Sun, C.-H., Li, W.-J., Yu, L.-Y., Zhou, J.-Q., Zhang, Yue-Qin, Xu, L.-H., Jiang, C.-L., 2007. Deinococcus yunweiensis sp. nov., a gamma– and UV-radiation-resistant bacterium from China. International Journal of Systematic and Evolutionary Microbiology 57, 370–375. 10.1099/ijs.0.64292-0

55. Zhang, Z., Zhang, Q., Lu, T., Zhang, J., Sun, L., Hu, B., Hu, J., Peñuelas, J., Zhu, L., Qian, H., 2022. Residual chlorine disrupts the microbial communities and spreads antibiotic resistance in freshwater. Journal of Hazardous Materials 423, 127152. 10.1016/j.jhazmat.2021.127152

56. Zhao, R., Symonds, J.E., Walker, S.P., Steiner, K., Carter, C.G., Bowman, J.P., Nowak, B.F., 2020. Salinity and fish age affect the gut microbiota of farmed Chinook salmon (*Oncorhynchus tshawytscha*). Aquaculture 528, 735539. 10.1016/j.aquaculture.2020.735539

57. Zhou, Q., Chen, W., Shan, K., Zheng, L., Song, L., 2014. Influence of sunlight on the proliferation of cyanobacterial blooms and its potential applications in Lake Taihu, China. Journal of Environmental Sciences 26, 626–635. 10.1016/S1001-0742(13)60457-X

58. Zhou, W., Li, W., Chen, J., Zhou, Y., Wei, Z., Gong, L., 2021. Microbial diversity in full-scale water supply systems through sequencing technology: a review. RSC Adv. 11, 25484– 25496. 10.1039/D1RA03680G

59. Zhu, H., Qiang, J., Li, Q., Nie, Z., Gao, J., Sun, Y., Xu, G., 2022. Multi-kingdom microbiota and functions changes associated with culture mode in genetically improved farmed tilapia (Oreochromis niloticus). Frontiers in Physiology 13.

