## Supplementary methods for "Microbiome Dynamics in Tank- and Pond-Reared GIFT Tilapia"

***Ectoparasite Assessment***

The density of ectoparasites, including dactylogyrus, trichodina, and ciliates, was assessed from individual skin, gill, and fin samples collected from all the tilapia used in this study, following a standardized method. In brief, skin samples consisted of scraped mucus and epithelial cells collected behind the pectoral fins and along the belly. For the gill and fin samples, small segments of the tips from a few gill filaments and a small portion of the pectoral fin were collected, respectively, using a tweezer and scissors. All samples were prepared, and parasite density was counted under a light compound microscope following the quick wet mount sampling guide for ectoparasites (WorldFish wet mount sampling guide (for ectoparasites & fungi) <https://digitalarchive.worldfishcenter.org/handle/20.500.12348/4837>.
