## Supplemental_Data3_PCA for "Microbiome Dynamics in Tank- and Pond-Reared GIFT Tilapia": Supplemental_Data3_Figure_PCA.pptx

### Slide 1
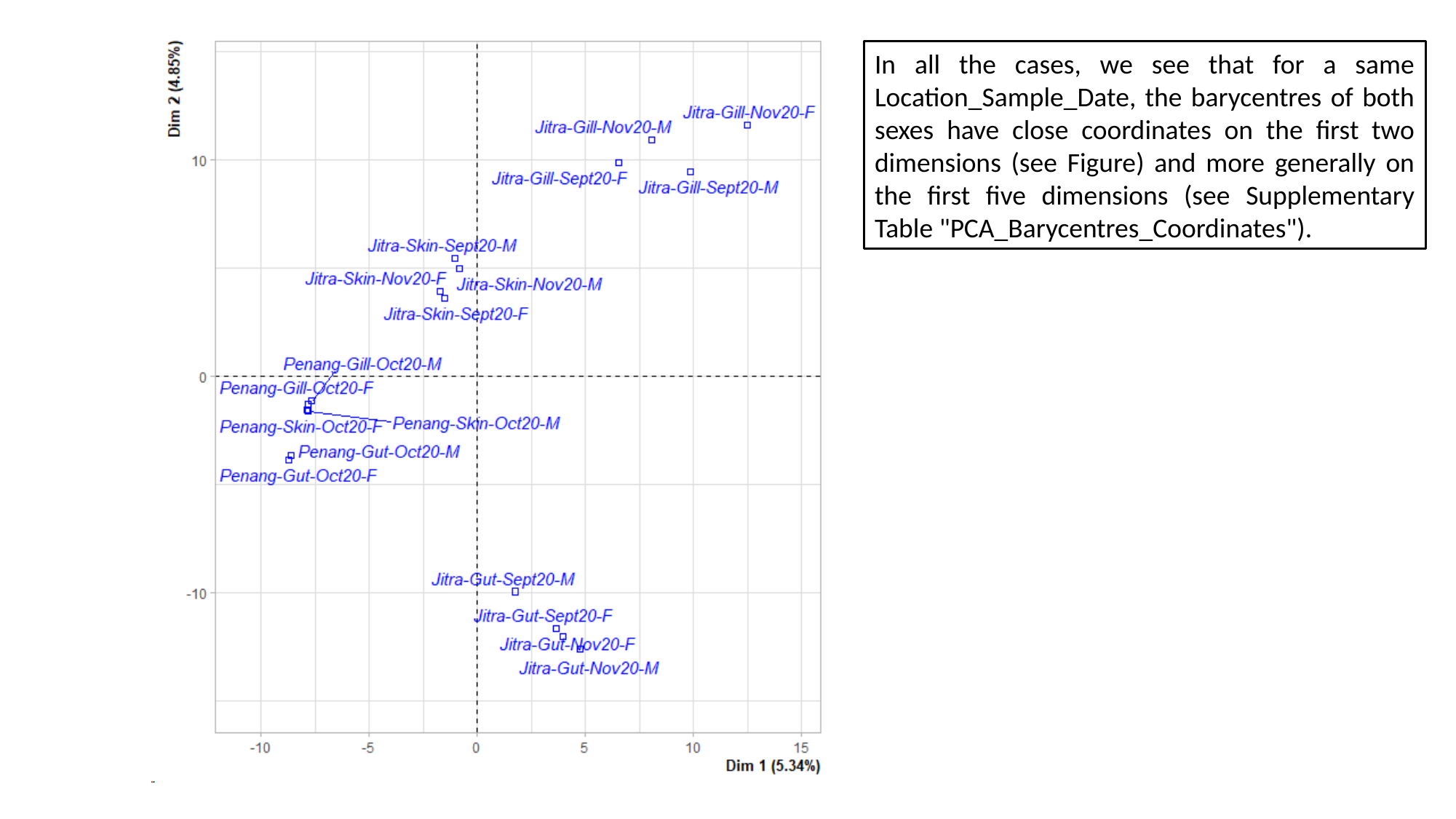

In all the cases, we see that for a same Location_Sample_Date, the barycentres of both sexes have close coordinates on the first two dimensions (see Figure) and more generally on the first five dimensions (see Supplementary Table "PCA_Barycentres_Coordinates").
